# Alternative mutational architectures producing identical M-matrices can lead to different patterns of evolutionary divergence

**DOI:** 10.1101/2023.08.11.553044

**Authors:** Daohan Jiang, Matt Pennell

## Abstract

Explaining macroevolutionary divergence in light of population genetics requires understanding the extent to which the patterns of mutational input contribute to long-term trends. In the context of quantitative traits, mutational input is typically described by the mutational variance-covariance matrix, or the **M**-matrix, which summarizes phenotypic variances and covariances introduced by new mutations per generation. However, as a summary statistic, the **M**-matrix does not fully capture all the relevant information from the underlying mutational architecture, and there exist infinitely many possible underlying mutational architectures that give rise to the same **M**-matrix. Using individual-based simulations, we demonstrate mutational architectures that produce the same **M**-matrix can lead to different levels of constraint on evolution and result in difference in within-population genetic variance, between-population divergence, and rate of adaptation. In particular, the rate of adaptation and that of neutral evolution are both reduced when a greater proportion of loci are pleiotropic. Our results reveal that aspects of mutational input not reflected by the **M**-matrix can have a profound impact on long-term evolution, and suggest it is important to take them into account in order to connect patterns of long-term phenotypic evolution to underlying microevolutionary mechanisms.

## Introduction

How structure of mutational input is constraining availability of standing genetic variation and ultimately shaping the course of long-term phenotypic evolution has been a question of great interest [Gould, 1980, Nei, 2013, Stoltzfus, 2021], and addressing this problem requires understanding the degree and pattern of mutational input. In studies of quantitative traits, abundance of mutational input is usually quantified using the mutational variance, defined as phenotypic variance introduced by new mutations per unit time, usually presented on a per-generation basis. For multi-dimensional traits, the mutational variance-covariance matrix (the **M**-matrix, hereafter **M**) is used to summarize the amount and correlational structure of mutational input simultaneously. Each diagonal element of **M** represents a trait’s mutational variance, and each off-diagonal element represents the mutational covariance (i.e., phenotypic covariance introduced by new mutations per generation) between two traits. To estimate **M**, one can use mutagenesis or mutation accumulation (MA) experiments to generate a large number of mutant genotypes and compute phenotypic (co)variances among them (e.g., [Camara and Pigliucci, 1999] and [Houle and Fierst, 2013]).

Often underappreciated is that **M** is only an insufficient summary statistic for the mutational architecture (i.e., the number of genomic loci affecting each trait, the mutation rate and spectrum at each locus, and the phenotypic effects of mutations). By definition, the mutational variance is a product of the mutation rate and variance of mutations’ effect on the trait; similarly, the mutational covariance of a given pair of traits is a product of the rate of pleiotropic mutations affecting both traits and the covariance of the mutations’ effects on two traits [Hansen, 2006, Lynch and Hill, 1986]. With each (co)variance being a product of different parameters, the same **M** can potentially result from different combination of parameters. Also underappreciated is how these different combinations could affect the evolutionary dynamics differently, which is also poorly understood.

As an illustration of the one-to-many mapping between **M** and the underlying mutational architectures, consider two quantitative traits, trait 1 (*z*_1_ hereafter) and trait 2 (*z*_2_ hereafter), and derive the mutational (co)variances from population genetic first principles. Genomic loci affecting these traits fall into three groups: there are *L*_1_ loci that exclusively affect *z*_1_, *L*_2_ loci that exclusively affect *z*_2_, and *L*_*P*_ loci that pleiotropically affect *z*_1_ and *z*_2_ simultaneously (*L*_1_, *L*_2_, and *L*_*P*_ are all non-negative integers). Let us assume each loci has two possible alleles, and all loci’s phenotypic effects are additive. Mutational variance resulting from loci that exclusively affect *z*_1_ is then given by [Lynch and Hill, 1986]

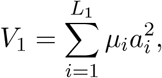

where *µ*_*i*_ is mutation rate of the *i*-th locus and *a*_*i*_ is the phenotypic effect of a mutation at the *i*-th locus. Similarly, mutational variance resulting from *z*_2_ is given by

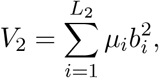

where *b*_*i*_ is the phenotypic effect of a mutation at the *i*-th locus.

Let us denote the effect of a mutation at the *i*-th pleiotropic locus as a vector *δz* = (*a*_*i*_, *b*_*i*_). The total mutational (co)variance contributed by pleiotropic locus is then given by

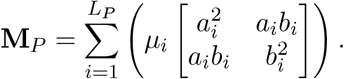

The mutational covariance matrix, or **M**-matrix for *z*_1_ and *z*_2_ is a sum of contribution from three types of loci:

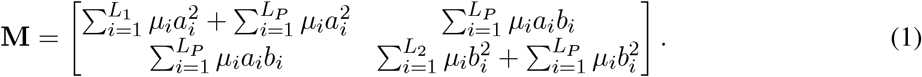

If all loci have the same mutation rate *µ*, the above equation becomes

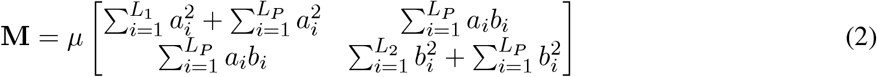

and if we also assume that mutation’s effect on a trait is normally distributed across loci, there is

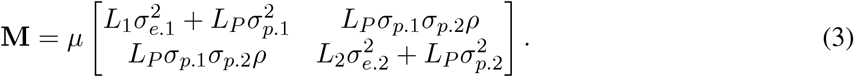

In the above equation, *σ*_*e*.1_ and *σ*_*e*.2_ are the standard deviations of phenotypic effects of mutations at loci that exclusively affect *z*_1_ and those at loci that exclusively affect *z*_2_, respectively. Standard deviations of pleiotropic mutations’ effects on *z*_1_ and *z*_2_ are *σ*_*p*.1_ and *σ*_*p*.2_, respectively. At last, *ρ* is the correlation coefficient between pleiotropic mutations’ effects on two traits. It can be seen that every element of **M** is a product of multiple quantities, and it is plausible that different combinations of them give rise to the same **M**. Below we will demonstrate how **M** can remain unchanged with multiple parameters in Eqn. 3 are altered. We denote to a particular vector of values ℙ, where ℙ: {*L*_1_, *σ*_*e*.1_, *L*_2_, *σ*_*e*.2_, *L*_*P*_, *ρ, σ*_*p*.1_, *σ*_*p*.2_} hereafter for convenience.

To see how we can manipulate the parameters while holding **M** constant, let *L*_*P*_, *ρ, σ*_*p*.1_, and *σ*_*p*.2_ each be multiplied by a rescaling coefficient, such that they become *C*_*P*_ *L*_*P*_, *C*_*ρ*_*ρ, C*_*p*.1_*σ*_*p*.1_, and *C*_*p*.2_*σ*_*p*.2_, respectively, where *C*_*P*_ *C*_*ρ*_*C*_*p*.1_*C*_*p*.2_ = 1 and *C*_*ρ*_ *<* 1*/* |*ρ*|. Let us multiply 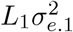 and 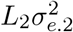 by rescaling coefficients *C*_1_ and *C*_2_, respectively, to keep the mutational variances unchanged:

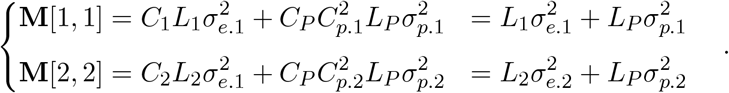

Solving the above equations gives

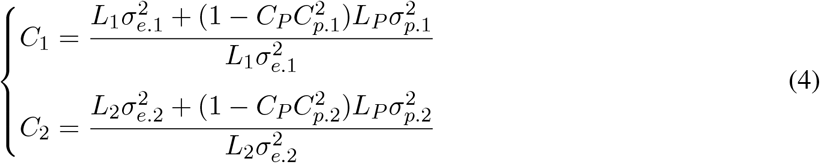

*C*_1_ and *C*_2_ must be non-negative as no mutation rate or standard deviation can be negative. Therefore, *C*_1_ and *C*_2_ can only be solved if

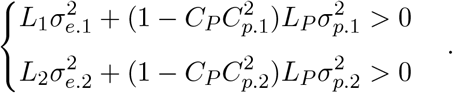

Solving the above system of inequalities gives

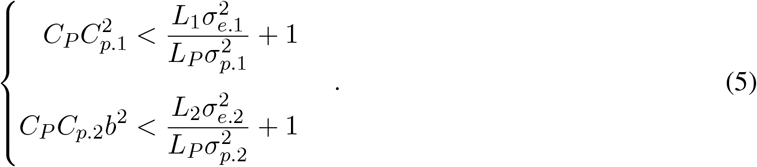

Hence, given **M**, certain combinations of *C*_*P*_, *C*_*ρ*_, *C*_*p*.1_, and *C*_*p*.2_ are guaranteed to alter the mutational variances. Biologically, if the portion of mutational variance attributable to pleiotropic mutations gets too high, it would be impossible to keep the total mutational variance unchanged by reducing the portion contributed by non-pleiotropic mutations. Given that *C*_1_ can be solved, the change to 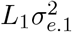 can be done by altering *L*_1_, *σ*_*e*.1_, or both. Thus, for any given combination of *C*_*P*_, *C*_*ρ*_, *C*_*p*.1_, and *C*_*p*.2_, there exists infinitely many ways to adjust 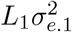 to keep **M** unchanged. Similarly, there are also infinitely many ways to adjust 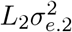. Hence, there exists infinitely many unique P that give rise to the same **M**.

In this study, we use population genetic simulations to explore dynamics of phenotypic evolution in the face of the same **M** but different underlying mutational architectures. Specifically, we examined series of scenarios where the fraction of loci that are pleiotropic varied, and show that both neutral evolution and adaptation are more constrained when the fraction is higher.

## Results and Discussion

To demonstrate how mutational architectures that produce identical **M**-matrices can lead to different evolutionary dynamics, we performed evolutionary simulations in SLiM [Haller and Messer, 2023] and examined phenotypic variation within and between populations at the end of the simulations. We considered genotype-phenotype (G-P) maps where each trait is affected by 50 genomic loci with equal effect size. Some loci are non-pleiotropic, whereas others are pleiotropic loci that affect all the traits. Different G-P maps being compared have different numbers of pleiotropic and non-pleiotropic loci, but the number of loci affecting each trait is constant (see Fig. 1 for a schematic illustration). Pleiotropic mutation’s effects on different traits are uncorrelated. Together, all these G-P maps produce the same mutational variances and zero mutational covariance (the **M**-matrices are identical).

**Figure 1:**
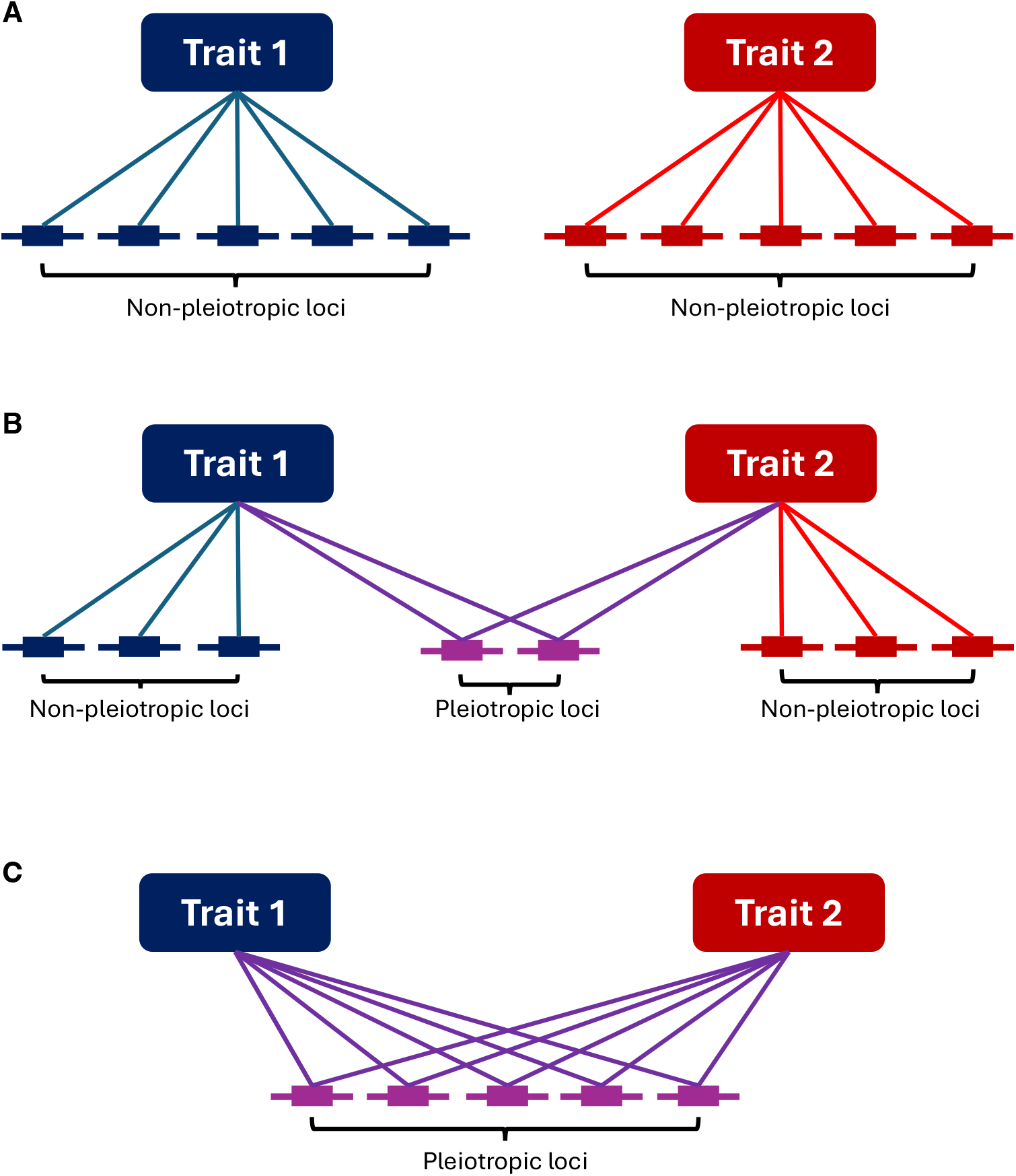
Schematic illustration of alternative genotype-phenotype maps that produce the same **M**-matrix. A locus’s effect on a trait is indicated by a line connecting the trait and the locus. In all three scenarios, each trait is affected by 5 loci, the distribution of mutations’ per-trait effect is the same for all loci, and pleiotropic mutation’s effect on two traits are uncorrelated. Thus, the two traits have the same mutational variance and zero genetic covariance in all scenarios. (A) Each trait affected by 5 non-pleiotropic loci. (B) Each trait is affected by 3 non-pleiotropic loci and 2 pleiotropic loci. (C) Both traits are affected by the same 5 loci.

We first examined scenarios where traits under concern are all under stabilizing selection. For each G-P map, we simulated 50 replicate populations, and examined within-population genetic variance (*V*_*G*_) and between-population variance (*V*_*R*_) at the end of simulation. While the different G-P maps showed little difference when only 2 traits were simulated, both *V*_*G*_ and *V*_*R*_ become lower when all loci are pleiotropic and each loci affects 5 or 10 traits (Fig. 2).

**Figure 2:**
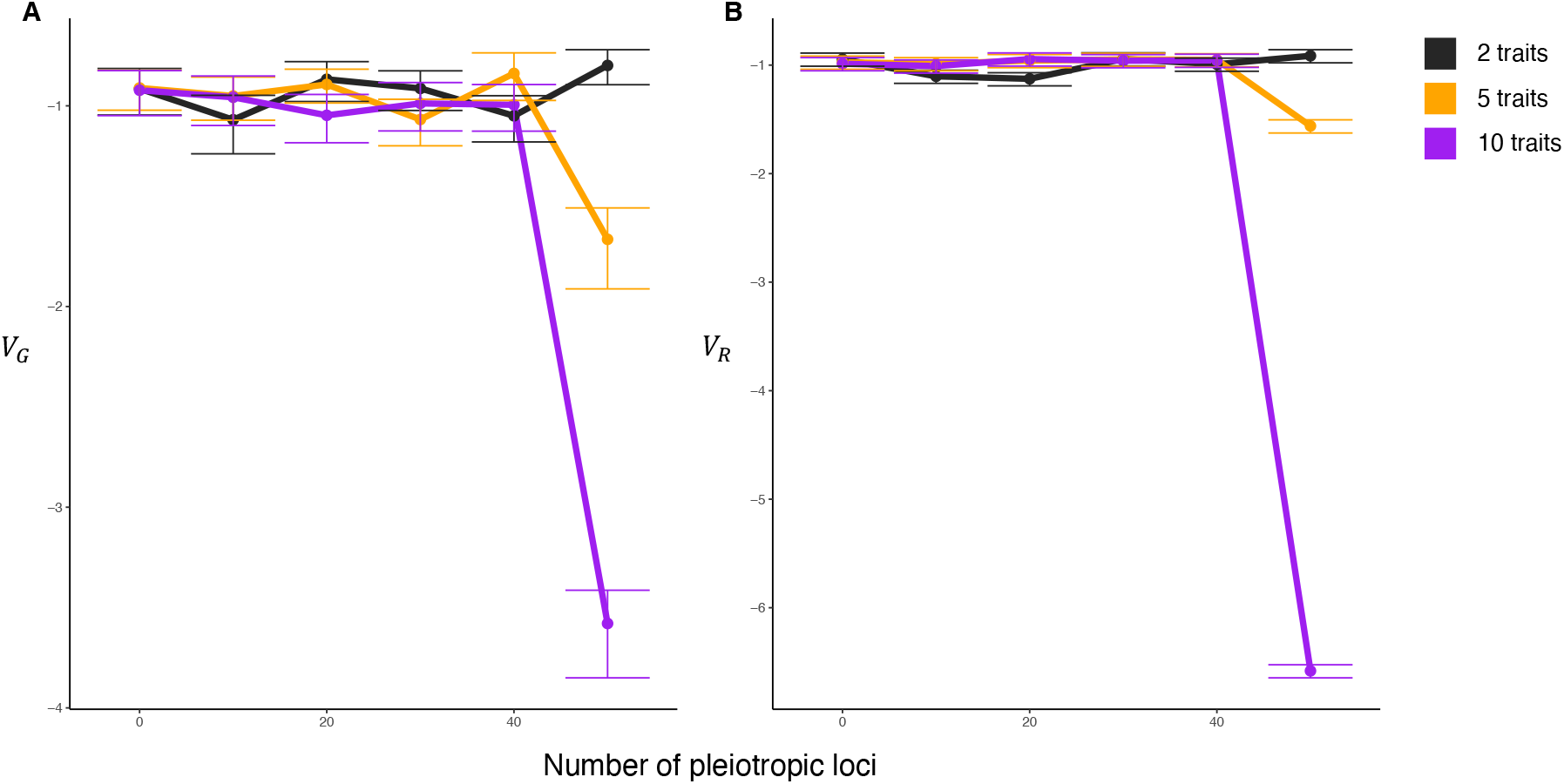
Phenotypic variance within and between populations when all traits are under stabilizing selection. Colors correspond to the number of traits being simulated. (A) Within-population genetic variance (*V*_*G*_), which is averaged across populations for each trait and then averaged across traits. Error bars reflect standard error, which is first calculated for each trait and then averaged across traits. (B) Between-population variance (*V*_*R*_), which is first calculated for each trait and then averaged across traits. Error bars reflect sampling standard deviation of sample variance at sample size of 50. Y-axes are in log10 scale.

We also examined the evolution of a neutral trait (i.e., *z*_1_) that does not affect fitness directly and asked how its evolution would be constrained by the indirect effect of other traits being under stabilizing selection. We predicted that, as the proportion of underlying loci of *z*_1_ increases, *V*_*G*_ and *V*_*R*_ of *z*_1_ will decrease. Indeed, when all loci are pleiotropic and each locus affects 10 traits, *V*_*G*_ and *V*_*R*_ of *z*_1_ both become magnitudes lower than those in other scenarios (Fig. 3). While *V*_*G*_ did not show clear trends when the level of pleiotropy is intermediate (i.e., not all loci are pleiotropic, the number of traits affected by each loci is relatively small), *V*_*R*_ decreased as the proportion of loci that are pleiotropic increased from 0 to 100% in scenarios of 5 and 10 traits (Fig. 3B). Note that even in the absence of pleiotropy, *V*_*R*_ of *z*_1_ is lower than the neutral expectation and lower when more traits are under stabilizing selection (Fig. 3B), indicating the rate of fixation of neutral mutations (i.e., non-pleiotropic mutations that affect *z*_1_ only) was reduced by unlinked background selection [Charlesworth, 2012, Matheson and Masel, 2024]. Together, our results show that prevalent pleiotropy can constrain the rate of neutral evolution as captured by phenotypic variance among lineages.

**Figure 3:**
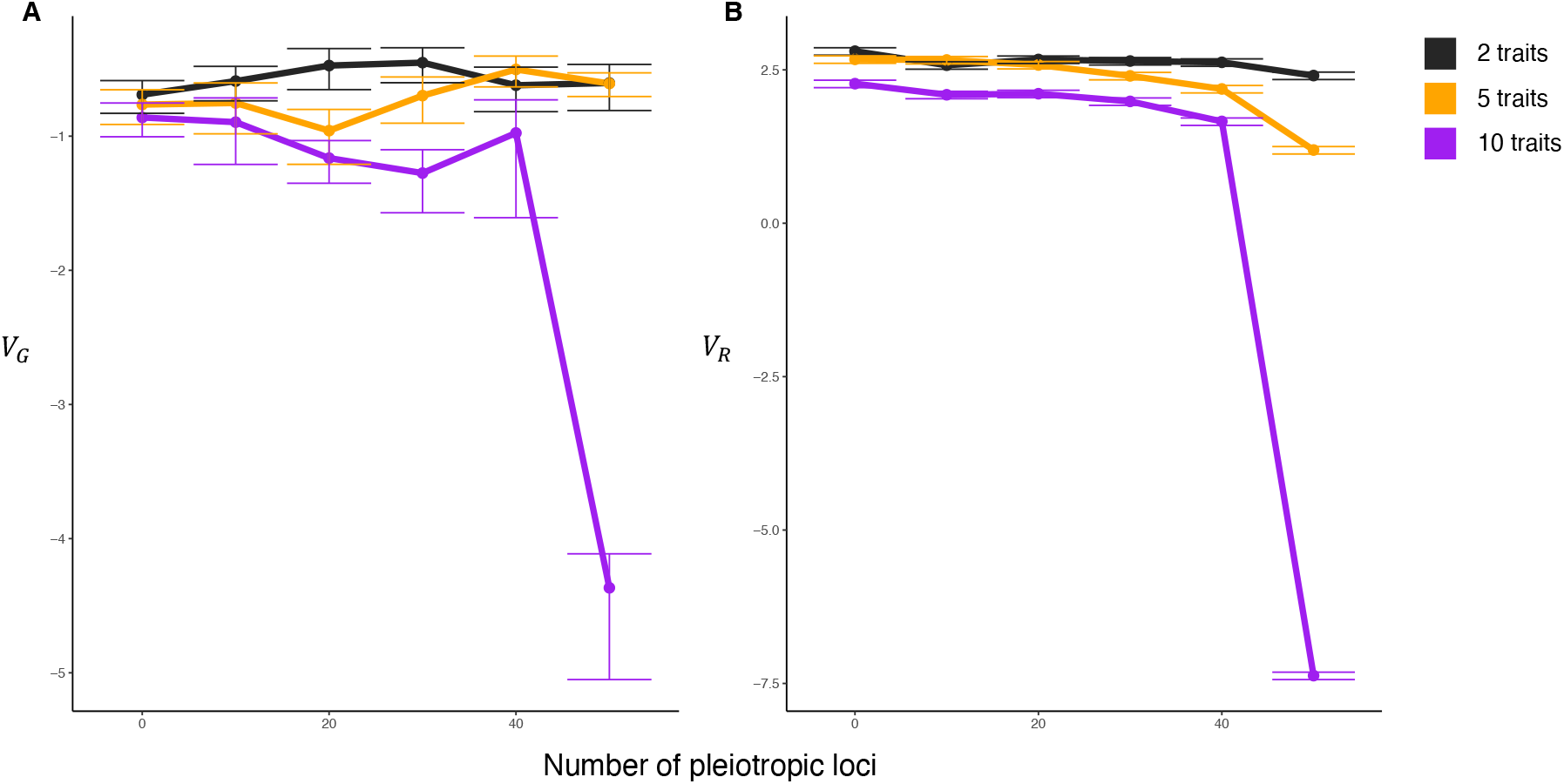
Variance of a neutral trait (*z*_1_) within and between populations when all other traits are under stabilizing selection. Colors correspond to the number of traits being simulated. (A) Within-population genetic variance (*V*_*G*_) of *z*_1_, which is averaged across populations. Error bars reflect standard error, which is first calculated for each trait and then averaged across traits. (B) Between-population variance (*V*_*R*_) of *z*_1_. Error bars reflect sampling standard deviation of sample variance at sample size of 50. Y-axes are in log10 scale.

And last, we asked how these different G-P maps could constrain adaptation when a specific trait (i.e., *z*_1_) is under directional selection and other traits are under stabilizing selection. Under such regimes of selection, selection on different traits can interfere, and pleiotropy can have a profound impact on a trait’s response to directional selection [Hansen and Houle, 2008]. We simulated evolution in non-Wright-Fisher (non-WF) populations whose size can change over time and examined their mean phenotypes and population sizes at the end of the simulations. Under our simulations’ conditions, an individual’s phenotype affects its viability while fecundity is invariable among individuals. As the population undergoes adaptive evolution, it will be able to reach and maintain a greater size as death rate is lower; when the population is well adapted (i.e., all individuals have the optimal phenotype), its size will stay close to the carrying capacity *K*, which is an upper limit to population imposed by the environmental condition. As pleiotropic loci are more likely to have detrimental effects on traits under stabilizing selection, the supply of adaptive mutations will be more limited when a greater fraction of loci are pleiotropic (Fig. S1), which could result in lower rate of adaptation and smaller population size. While it is not impossible for a population with very low rate of adaptation to reach the optimum in the end if it is given unlimited time [Sella, 2009], actual populations do not evolve in constant environments indefinitely, and it is often the dynamics of adaptation during a transient period rather than the long-term equilibrium in a static environment that is most relevant (e.g., in the context of evolutionary rescue [Anciaux et al., 2018, Orr and Unckless, 2014] or fluctuating selection [Holstad et al., 2024]). Thus, we let the simulation run for a fixed amount of time, and examined the evolved populations’ sizes and mean phenotypes at the end. As predicted, as the proportion of loci that are pleiotropic increased, population size at the end decreased (Fig. 4A) and the population mean of 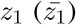 became farther away from the optimum (Fig. 4B). When the number of pleiotropic loci is no more than 20 (i.e., 40% of loci underlying each trait), population size at the end was close to *K*, and 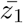 was close to the optimum, indicating successful adaptation. In contrast, when all loci are pleiotropic and the number of traits affected by each locus is large (i.e., 5 or 10 traits), many populations underwent no adaptive evolutionary change at all within time of simulation (Table S1).

**Figure 4:**
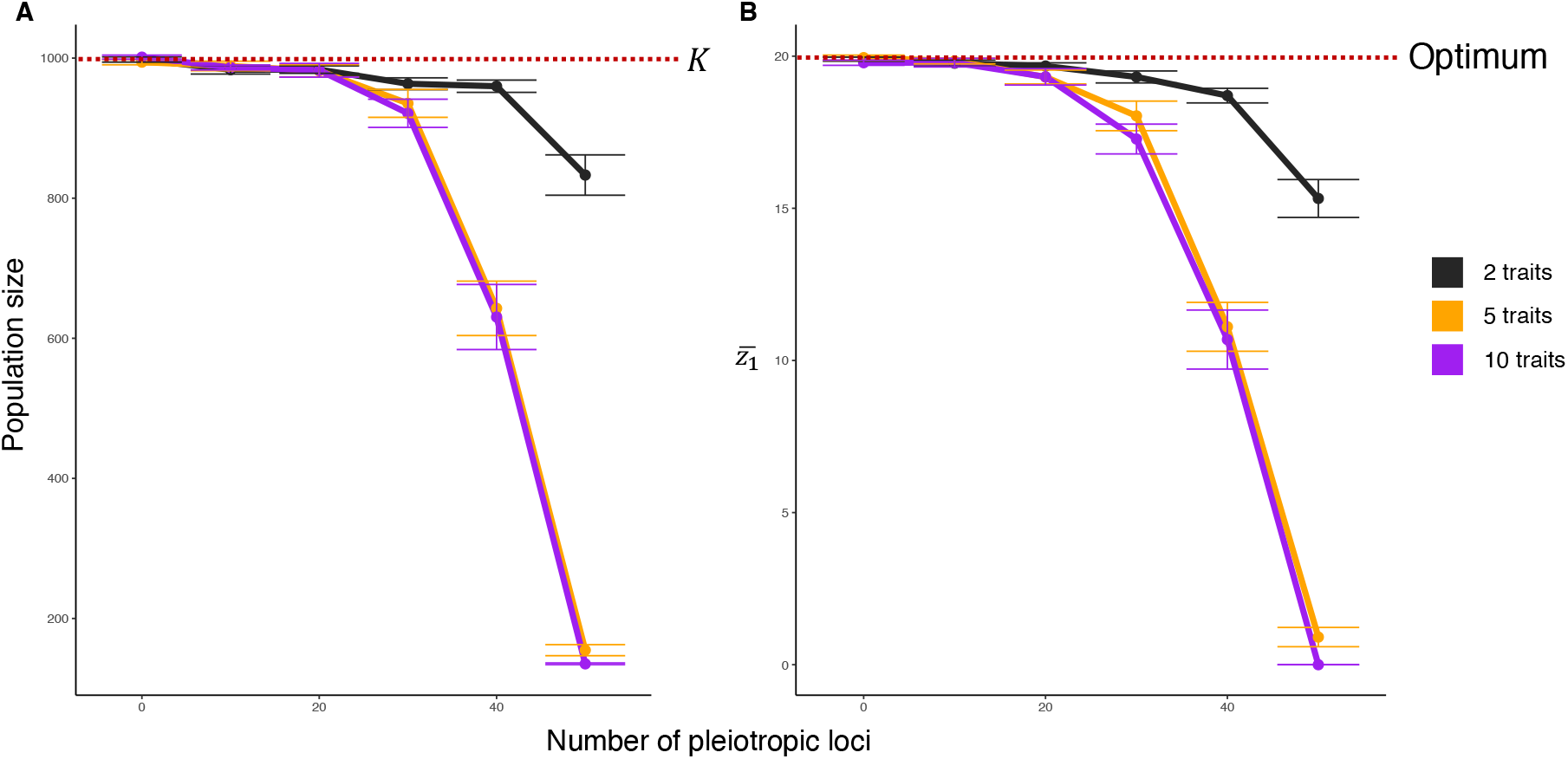
Adaptive evolution in non-Wright-Fisher populations. Colors correspond to the number of traits being simulated. (A) Mean population size at the end of simulation. Red dashed line represents the carrying capacity (*K*). (B) Mean value of trait under directional selection 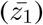 at the end of simulation. Red dashed line represents its optimum. Error bars in both panels reflect standard error.

**Table 1:** Definition of simulation parameters.

| Parameter | Definition |
| --- | --- |
| $z_1, z_2$ | Phenotypic trait 1 and trait 2. |
| $\mathbf{M}$ | Mutational variance-covariance matrix, **M**-matrix. |
| $V_1, V_2$ | Mutational variances of $z_1$ and $z_2$, respectively. |
| $L_1$ | The number of loci that exclusively affect $z_1$. |
| $L_2$ | The number of loci that exclusively affect $z_2$. |
| $L_P$ | The number of pleiotropic loci that affect both $z_1$ and $z_2$. |
| $\mu$ | Per-locus mutation rate; $\mu_i$ denotes mutation rate of the $i$-th locus. |
| $a_i, b_i$ | Effects of a mutation at the $i$-th locus on $z_1$ and $z_2$, respectively. |
| $\sigma_{e.1}$ | Standard deviations of phenotypic effect of mutations that exclusively affect $z_1$. |
| $\sigma_{e.2}$ | Standard deviations of phenotypic effect of mutations that exclusively affect $z_2$. |
| $\sigma_{p.1}, \sigma_{p.2}$ | Standard deviations of pleiotropic mutations' effect on $z_1$ and $z_2$, respectively. |
| $\rho$ | Correlation between pleiotropic mutations' effect on $z_1$ and $z_2$. |
| $\omega$ | Fitness of an individual with respect to traits under concern. |
| $\mathbf{S}$ | Matrix characterizing multivariate selection. |
| $N$ | Size of a population; $N_t$ denotes population size at time $t$. |
| $K$ | Carrying capacity; equilibrium population size when the phenotype is optimized. |

Together, our simulation results show mutational architectures that produce the same **M**-matrix but have distinct “hidden” properties can have drastically different effects on dynamics of neutral phenotypic evolution and adaptation. The effect of hidden aspects of the mutational architecture on phenotypic evolution has important implications for understanding mechanisms of phenotypic evolution in nature. That mutational input constrains availability of genetic variance and ultimately long-term phenotypic evolution is a long-standing and controversial hypothesis [Gould, 1980, Nei, 2013, Stoltzfus, 2021]; in principle, one can test it by comparing **M** to patterns of within-species additive genetic (co)variances (as encapsulated by the genetic variance-covariance matrix, **G**) and evolutionary (co)variance among species (as encapsulated by the evolutionary variance-covariance matrix, **R**) (e.g., [Houle et al., 2017]); strong similarity between **M** and the other two matrices would be consistent with the patterns of mutational input driving long-term evolution. However, this test faces conceptual difficulties and is not as straightforward as it appears to be: as the dispositional effect of mutational input on evolution cannot be learned from the **M**-matrix alone, a comparison of matrices alone is also not sufficient to tell whether and how mutational constraints have shaped observed phenotypic divergence.

The key difference between mutational architectures examined in this study is in their degree of pleiotropy, specifically the proportion of loci that are pleiotropic along underlying loci of each trait. We found that pleiotropic mutations are generally more deleterious, less likely to be adaptive, and less likely to fix, resulting in constraints on both neutral and adaptive evolution. Our findings regarding the effect of pleiotropy on evolution agree with those of earlier studies [Battlay et al., 2024, Chevin et al., 2010, Jiang and Zhang, 2020, Martin, 2014, McGuigan, 2006, Orr, 2000], but further show that this effect persists even given the same **M**. If the effect of details of pleiotropy is overlooked and assumed to make little difference to evolution, conclusions about phenotypic evolution that are contingent on strong assumptions about pleiotropy could be mis-interpreted as general. In particular, models for the evolution of multivariate traits often assume universal pleiotropy (i.e., every mutation affects every trait), which can have substantial impact on their conclusions and implications. For instance, Fisher’s Geometric Model (FGM) makes this assumption, which leads to the prediction that mutations with smaller effect sizes are more likely to be adaptive and that there is a “cost of complexity” as adaptation is slower when there are a greater number of phenotypic dimensions [Fisher, 1930, Orr, 2000, Tenaillon, 2014, Welch and Waxman, 2003]. Similarly, in a series of modeling studies, Jones et al. [2007] and Jones et al. [2014] assumed universal pleiotropy when modeling the evolution of the mutational architecture under second-order selection, making the effect size correlation being the only evolvable aspect of mutational architecture; it is unknown whether the mutational architecture would evolve differently if the assumption of universal pleiotropy is relaxed. The degree to which the assumption of universal pleiotropy is reasonable remains an open question [Boyle et al., 2017, Hill and Zhang, 2012a,b, Paaby and Rockman, 2013, Wagner and Zhang, 2011, Zhang, 2023]. Some studies have found that each gene or mutation typically affects only a small subset of traits and suggested that adaptation is not necessarily more constrained in complex organisms as FGM would indicate [Ho and Zhang, 2014, Wagner et al., 2008, Wang et al., 2010]. Others argue that pleiotropy is more pervasive and that many empirical studies underestimate the prevalence of pleiotropy due to technical issues [Hill and Zhang, 2012b]. Furthermore, the recently proposed “omnigenic” model [Boyle et al., 2017, Liu et al., 2019] argues that, because of properties of the regulatory network, each individual gene or mutation can affect a large number of traits while having major effects on a small number of traits. No matter how the debate would resolve, it is clear we cannot take the universal pleiotropy assumption for granted, and it is essential for future studies to be cautious when modeling the evolution multivariate traits and interpreting observed phenotypic variations. It is also worth noting that pleiotropy makes a difference even when mutations’ effects on different traits are uncorrelated. Correlated pleiotropic effects, which manifest as mutational covariances, are known to shape the structure of genetic covariances and eventually patterns of correlated evolution [Lande, 1979, 1980, Wagner, 1989] whereas the effect of unstructured pleiotropy on evolution is less appreciated. Nevertheless, unstructured pleiotropy can alter the distribution of effects of new mutations, potentially constraining the course of evolution. Together, we suggest that, with only the **M**-matrix along with regime of selection, robust predictions about the course of evolution cannot be made without further information, and more detailed understanding of the mutational architecture would be essential for understanding mechanisms of phenotypic evolution.

## Conclusion

In this study, we show that the **M**-matrix, a summary statistic commonly used to describe mutational input for quantitative traits, does not fully capture key features of the mutational architecture even when mutations’ effects are all additive. Using simulations, we show difference in properties of these mutational architectures can result in different evolutionary dynamics. Specifically, when a greater fraction of loci affecting a given trait are pleiotropic, the trait under concern will have lower rates of neutral evolution and adaptation. We suggest that hidden aspects of mutational architectures that are not reflected by **M**-matrices poses significant challenge to attempts to understand mechanisms of phenotypic evolution and requires more explicit consideration in future studies.

## Methods

### Genotype-phenotype maps

We considered a set of quantitative traits, each affected by a set of underlying loci (i.e., genes or genomic regions). We considered an infinite sites model where mutations at any given locus are all distinct from each other and recurrent mutations never occur. Therefore, in our simulations, each mutation’s phenotypic effect is sampled independently from the locus-specific distribution. Effects of mutations on each trait were additive. For simplicity, heritability was assumed to be 100% for all traits. Two types of loci were considered in our simulations: non-pleiotropic loci that each affects a single trait, and universally pleiotropic loci that affect all traits. When a mutation occurs at a non-pleiotropic locus, its effect on the trait to be affected was sampled from a normal distribution 𝒩 (0, *σ*); in our simulations, we had *σ* = 1 for all non-pleiotropic loci. If a mutation occurs at a pleiotropic locus, its effect is sampled from a multivariate distribution characterized by an identity matrix. We assumed no bias in mutation’s phenotypic effect; that is, the mean effect of mutations at any given locus on any given trait was zero. We let every trait under consideration have 50 underlying loci, and compared G-P maps where 0, 10, 20, 30, 40, and 50 of these loci are pleiotropic. We considered scenarios where 2, 5, and 10 traits are affected by each pleiotropic locus. Note that in the case of no pleiotropy, we also performed simulations with 2, 5, and 10 traits.

### Selection on phenotypic traits

We considered a multivariate Gaussian fitness function, which is described by a covariance matrix **S**. Each diagonal element of **S** is the width of an individual trait’s fitness function (i.e., variance of a normal distribution), and off-diagonal elements represent correlational selection for relationships between traits [Arnold et al., 2008, 2001].

To calculate fitness given the *n*-dimensional phenotype 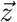, we first calculate its distance to the optimal phenotype 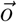:

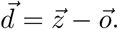

We then calculate the projection of 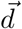 on eigenvectors of **S**:

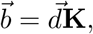

where **K** is the eigenvector matrix of **S**. Fitness is then calculated as

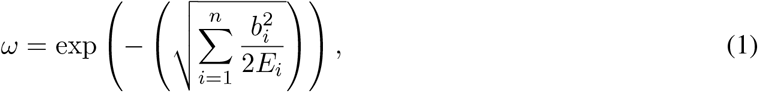

where *b*_*i*_ is the *i*-th element of 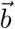 and *E*_*i*_ is the *n*-th eigenvalue of **S**. If an eigenvalue of **S** (e.g., *E*_*i*_) is zero, the corresponding term in Eqn. (1) 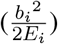 would be dropped. The biological interpretation of such a situation is the lack of selection on a specific phenotypic dimension, in which case the phenotypic dimension with no selection should not be considered when calculating fitness.

In our simulations, we only considered scenarios without correlational selection, so **S**-matrices being considered were all diagonal. Eqn. (1) thus becomes

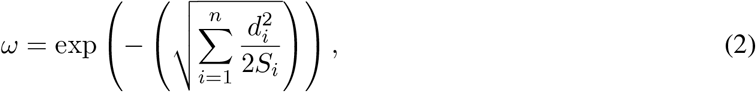

where *d*_*i*_ is the *i*-th element of 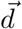 and *S*_*i*_ is the *i*-th diagonal element of **S**, characterizing strength of selection on the *i*-th trait.

We had all traits start from a value of 0 in our simulations. All traits’ optimal values are equal to 0, unless noted otherwise. Diagonal elements of **S** are all equal to 1, unless noted otherwise. In simulations where one trait (i.e., *z*_1_) is neutral, the corresponding diagonal element of **S**, *S*_1,1_ is equal to 0 and the trait is not counted when calculating fitness. In simulations where one trait (i.e., *z*_1_) is under directional selection, we set trait’s optimal value to be 20 and *S*_1,1_ = 100. Under such a setting, it requires multiple substitutions for the phenotype to be optimized and the initial fitness is not too low to cause quick extinction such that it is easy to quantify and visualize rate of adaptation using the population mean phenotype at the end.

### SLiM simulations

We simulated the evolution of orthogonal traits with zero mutational covariance in diploid, hermaphrodite, and free-mating populations in SLiM [Haller and Messer, 2023]. Each locus that affect trait(s) was represented as a single genetic element object in SLiM. Each locus’s mutation rate was set to be 10^−6^ per generation. We also assumed free recombination between loci and no recombination within loci (i.e., causal loci sparsely distributed along the chromosome). Fitness with respect to traits under consideration is calculated following Eqn. (2).

We simulated evolution of both Wright-Fisher (WF) and non-WF diploid populations. All WF populations had population size *N* = 1000, and simulation for each population lasted for 10*N* = 10^4^ ticks (i.e., generations). In the WF simulation, each individual’s fitness value is equal to fitness with respect to traits of concern. Simulation for each non-WF population started with *N* = *K* = 1000, where *K* is the carrying capacity, and ran for 10*K* = 10000 ticks. Reproduction takes place at the beginning of each tick, and the expected number of offspring produced by each individual each time was set to be 1, which was set to be the same for all individuals. Variation in fitness between individuals is mediated by death probability. The fitness value of a given individual (i.e., the *i*-th individual) at a given time *t* is calculated as 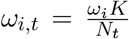, where *ω*_*i*_ is its fitness with respect to the traits under concern and *N*_*t*_ is the population size at the moment. If, after reproduction, an individual’s fitness is equal to or greater than 1, it will survive at the end of the tick; if all individuals’ fitness values are equal to or greater than 1, the population will grow.

For each evolutionary scenario, we simulated 50 replicate populations, which correspond to 50 subpopulation objects with zero gene flow in SLiM. Genetic variance (*V*_*G*_) of each trait was computed as phenotypic variance among individuals in a population at the end of the simulation. For each trait, genetic variances from the 50 replicate populations were averaged to represent the expected genetic variance. For scenarios where traits were either under stabilizing selection or no selection, we quantified the degree of evolutionary divergence among population using variance of mean phenotypes among replicate populations (*V*_*R*_). Because all traits under consideration had the same mutational variance, we averaged different traits’ *V*_*G*_ and *V*_*R*_ for simulation setting to represent the overall degree of constraint in the corresponding scenario. When a trait is under directional selection, we examined its mean across populations at the end; for non-WF simulations, population that had zero population sizes in the end where excluded calculating this mean phenotype.

## Code and data availability

Code and data files are available at https://github.com/phylo-lab-usc/m-matrix/tree/main.

## Acknowledgements

We thank Alex Cope, Joshua Schraiber, Arlin Stoltzfus, David McCandlish, and Joanna Masel for their thoughtful comments on this work. This study was supported by NIH grant R35GM151348 to MP.

## Supplementary materials

**Figure S1:**
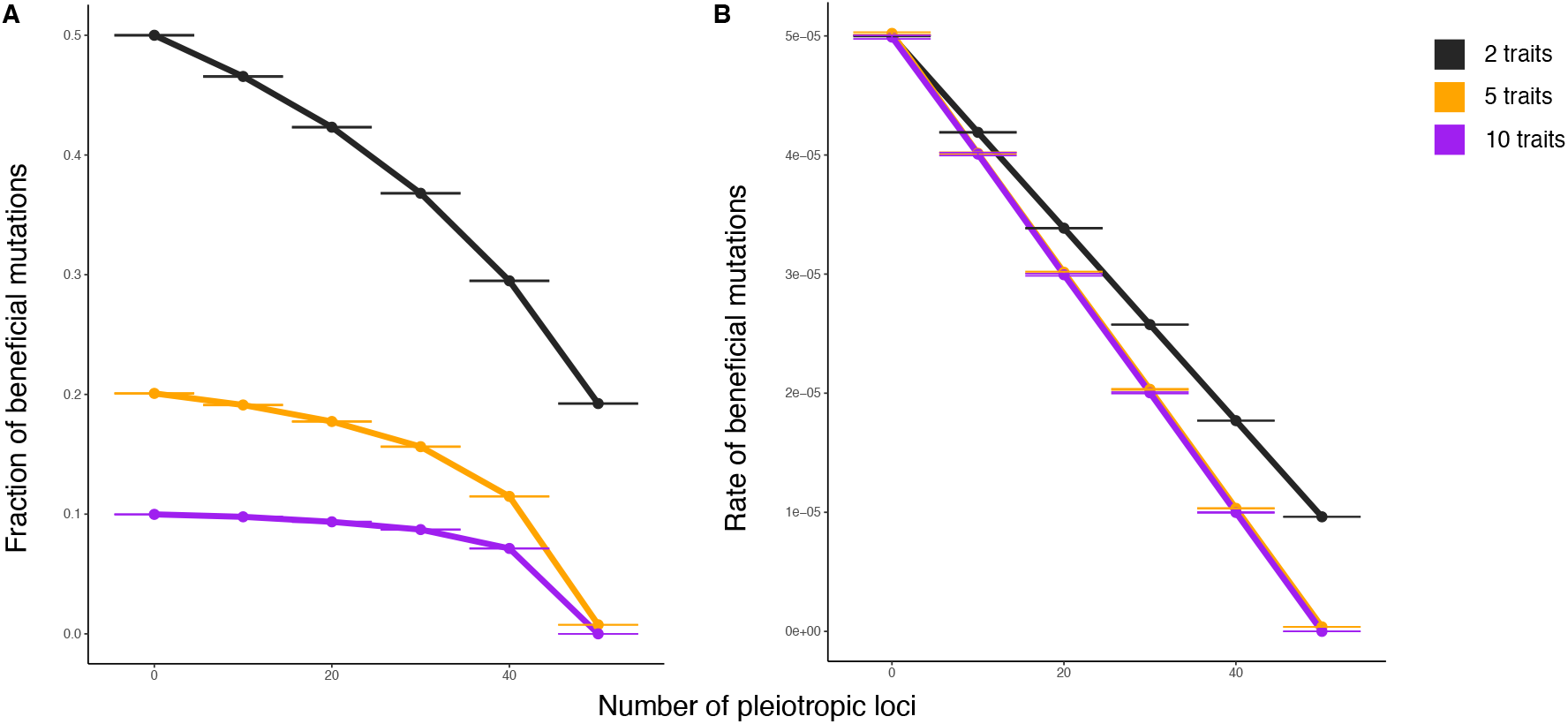
Frequency and rate of beneficial mutations when one trait is under directional selection and all other traits are under stabilizing selection. (A) Fraction of mutations that are beneficial estimated from 10^6^ random mutants. Error bars represent standard deviation of sample proportion at sample size of 10^6^. Fitness effect of each mutation is evaluated on the ancestral background at the beginning of the simulations. (B) Rate of beneficial mutations per genome per generation. Size of each error bar is equal to size of the corresponding error bar in (A) multiplied by the total mutation rate per genome per generation.

**Table S1:**
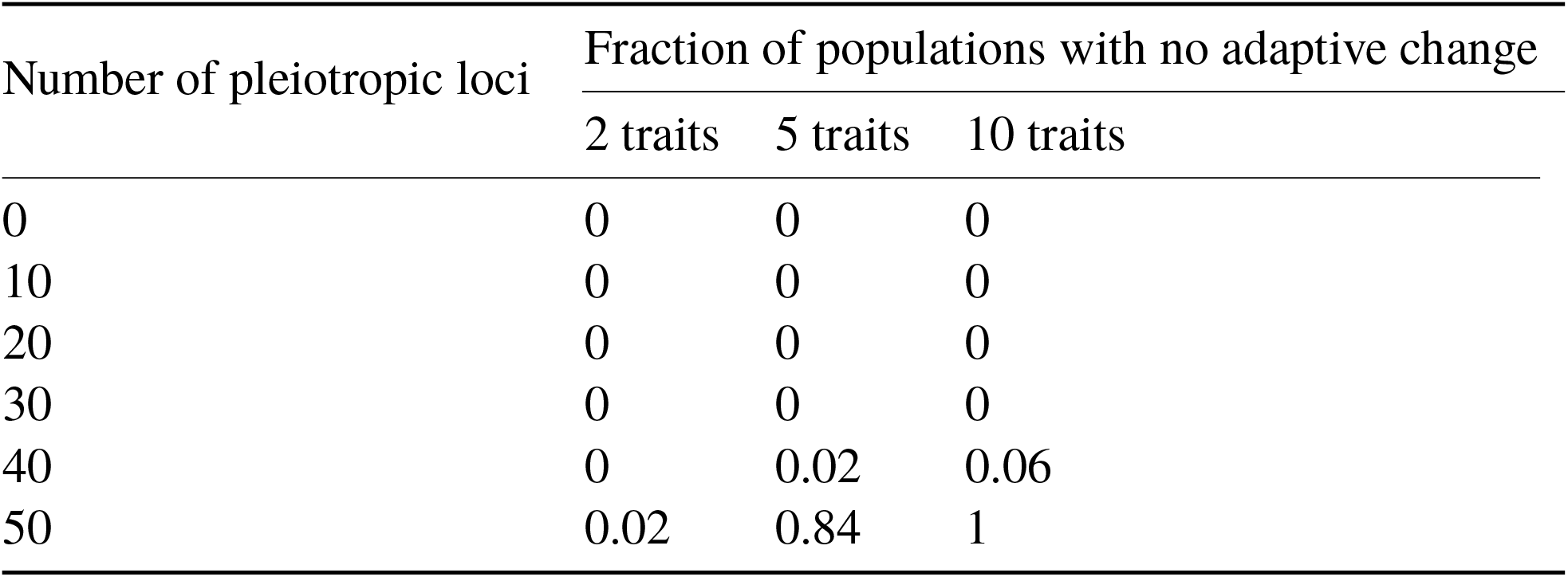
Fraction of replicate populations that underwent no adaptive evolutionary change (i.e., 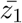 at the end of simulation is identical to that at the beginning).

